## Supplemental Table S1 for "Spatial distribution of ticks and tick-borne pathogens in central Hokkaido, Japan and associated ecological factors revealed by intensive short-term survey in 2024"

| Site No. | *Ixodes ovatus* adult | *Ixodes persulcatus* adult | *Ixodes pavlovskyi* adult | *Ixodes tanuki* adult | *Haemaphysalis megaspinosa* adult | *Haemaphysalis flava* adult | *Haemaphysalis japonica* adult | *Haemaphysalis longicornis* adult | *Haemaphysalis concinna* adult | *Ixodes* spp. nymph | *Haemaphysalis* spp. nymph | TBEV-positive sampe | YEZV-positive sampe | BJNV-positive sampe | LDB-positive sampe | pLDB-positive sample | RFB-positive sampe |
|---|---|---|---|---|---|---|---|---|---|---|---|---|---|---|---|---|---|
| 1 | 2 | 4 | 0 | 0 | 3 | 0 | 0 | 0 | 0 | 1 | 21 | 0 | 0 | 0 | 0 | 0 | 0 |
| 2 | 63 | 15 | 1 | 0 | 1 | 1 | 0 | 0 | 0 | 1 | 14 | 0 | 1 | 0 | 8 | 1 | 0 |
| 3 | 29 | 2 | 1 | 0 | 0 | 1 | 0 | 0 | 0 | 0 | 18 | 0 | 0 | 0 | 4 | 0 | 0 |
| 4 | 40 | 1 | 1 | 0 | 1 | 0 | 0 | 0 | 0 | 0 | 14 | 0 | 0 | 0 | 4 | 1 | 1 |
| 5 | 64 | 9 | 3 | 1 | 2 | 1 | 0 | 0 | 0 | 5 | 70 | 0 | 0 | 0 | 11 | 2 | 0 |
| 6 | 17 | 1 | 0 | 0 | 3 | 1 | 0 | 0 | 0 | 0 | 49 | 0 | 0 | 0 | 2 | 0 | 0 |
| 7 | 143 | 2 | 0 | 0 | 0 | 0 | 0 | 0 | 0 | 1 | 2 | 1 | 0 | 0 | 12 | 0 | 0 |
| 8 | 0 | 1 | 1 | 0 | 4 | 2 | 0 | 0 | 0 | 1 | 98 | 0 | 0 | 0 | 1 | 1 | 0 |
| 9 | 4 | 2 | 0 | 0 | 10 | 3 | 0 | 0 | 0 | 5 | 103 | 0 | 0 | 0 | 0 | 0 | 0 |
| 10 | 64 | 7 | 1 | 0 | 0 | 2 | 0 | 0 | 0 | 2 | 60 | 0 | 0 | 0 | 7 | 0 | 0 |
| 11 | 5 | 1 | 0 | 0 | 1 | 1 | 0 | 0 | 0 | 0 | 21 | 0 | 0 | 0 | 1 | 0 | 0 |
| 12 | 23 | 5 | 0 | 0 | 0 | 0 | 0 | 0 | 0 | 8 | 6 | 0 | 0 | 1 | 5 | 2 | 0 |
| 13 | 34 | 10 | 0 | 0 | 0 | 0 | 0 | 0 | 0 | 2 | 0 | 0 | 0 | 0 | 5 | 2 | 0 |
| 14 | 66 | 17 | 1 | 0 | 0 | 0 | 0 | 0 | 0 | 2 | 1 | 0 | 0 | 2 | 10 | 3 | 0 |
| 15 | 9 | 12 | 0 | 0 | 0 | 0 | 0 | 0 | 0 | 3 | 0 | 0 | 0 | 0 | 4 | 2 | 0 |
| 16 | 70 | 20 | 0 | 0 | 1 | 1 | 0 | 0 | 0 | 5 | 0 | 0 | 0 | 0 | 12 | 3 | 1 |
| 17 | 60 | 14 | 0 | 0 | 0 | 0 | 0 | 0 | 0 | 0 | 1 | 0 | 0 | 0 | 8 | 2 | 0 |
| 18 | 193 | 30 | 0 | 0 | 0 | 0 | 0 | 0 | 0 | 0 | 0 | 0 | 0 | 0 | 23 | 4 | 2 |
| 19 | 137 | 2 | 0 | 0 | 0 | 0 | 0 | 0 | 0 | 0 | 2 | 1 | 0 | 1 | 11 | 0 | 1 |
| 20 | 1 | 1 | 0 | 0 | 8 | 1 | 0 | 0 | 0 | 1 | 123 | 0 | 0 | 0 | 1 | 0 | 0 |
| 21 | 0 | 0 | 0 | 0 | 0 | 2 | 0 | 0 | 0 | 0 | 248 | 0 | 0 | 0 | 0 | 0 | 0 |
| 22 | 0 | 0 | 0 | 0 | 10 | 0 | 0 | 0 | 0 | 0 | 125 | 0 | 0 | 0 | 0 | 0 | 0 |
| 23 | 0 | 1 | 0 | 0 | 4 | 0 | 0 | 0 | 0 | 1 | 106 | 0 | 0 | 0 | 0 | 0 | 0 |
| 24 | 29 | 6 | 0 | 0 | 0 | 0 | 0 | 0 | 0 | 0 | 0 | 0 | 0 | 0 | 4 | 1 | 0 |
| 25 | 37 | 8 | 3 | 0 | 3 | 4 | 0 | 0 | 0 | 0 | 22 | 0 | 0 | 0 | 6 | 2 | 1 |
| 26 | 0 | 0 | 0 | 0 | 0 | 0 | 0 | 0 | 0 | 0 | 10 | 0 | 0 | 0 | 0 | 0 | 0 |
| 27 | 0 | 0 | 0 | 0 | 0 | 2 | 0 | 0 | 0 | 1 | 95 | 0 | 0 | 0 | 0 | 0 | 0 |
| 28 | 0 | 0 | 0 | 0 | 2 | 3 | 0 | 0 | 0 | 1 | 87 | 0 | 0 | 0 | 0 | 0 | 0 |
| 29 | 4 | 1 | 0 | 0 | 0 | 2 | 0 | 0 | 0 | 0 | 86 | 0 | 0 | 0 | 1 | 0 | 0 |
| 30 | 0 | 0 | 0 | 0 | 9 | 1 | 0 | 2 | 0 | 1 | 228 | 0 | 0 | 0 | 0 | 0 | 0 |
| 31 | 0 | 0 | 0 | 0 | 3 | 0 | 0 | 0 | 0 | 0 | 412 | 0 | 0 | 0 | 0 | 0 | 0 |
| 32 | 11 | 1 | 0 | 0 | 8 | 1 | 3 | 0 | 0 | 4 | 123 | 0 | 0 | 0 | 2 | 0 | 0 |
| 33 | 66 | 6 | 0 | 0 | 9 | 0 | 6 | 0 | 0 | 7 | 71 | 0 | 2 | 0 | 6 | 1 | 0 |
| 34 | 70 | 4 | 0 | 0 | 4 | 2 | 0 | 0 | 0 | 2 | 107 | 0 | 0 | 0 | 10 | 0 | 0 |
| 35 | 31 | 3 | 0 | 0 | 8 | 1 | 3 | 0 | 0 | 5 | 108 | 0 | 0 | 0 | 2 | 0 | 2 |
| 36 | 43 | 5 | 0 | 0 | 1 | 0 | 0 | 0 | 0 | 2 | 48 | 0 | 0 | 1 | 6 | 1 | 0 |
| 37 | 8 | 1 | 0 | 0 | 0 | 0 | 0 | 0 | 0 | 2 | 107 | 0 | 0 | 0 | 3 | 1 | 0 |
| 38 | 12 | 0 | 0 | 0 | 3 | 0 | 0 | 0 | 0 | 2 | 80 | 0 | 0 | 0 | 1 | 0 | 1 |
| 39 | 9 | 0 | 0 | 0 | 0 | 0 | 0 | 1 | 0 | 0 | 9 | 0 | 0 | 0 | 1 | 0 | 0 |
| 40 | 1 | 0 | 0 | 0 | 1 | 2 | 0 | 18 | 0 | 0 | 160 | 1 | 0 | 0 | 0 | 0 | 0 |
| 41 | 1 | 0 | 0 | 0 | 2 | 0 | 0 | 2 | 0 | 0 | 17 | 0 | 0 | 0 | 0 | 0 | 0 |
| 42 | 0 | 0 | 0 | 0 | 3 | 0 | 0 | 23 | 0 | 0 | 192 | 0 | 0 | 0 | 0 | 0 | 0 |
| 43 | 0 | 0 | 0 | 0 | 0 | 0 | 0 | 0 | 0 | 0 | 0 | 0 | 0 | 0 | 0 | 0 | 0 |
| 44 | 10 | 3 | 0 | 0 | 0 | 1 | 0 | 0 | 0 | 0 | 0 | 0 | 0 | 0 | 2 | 0 | 0 |
| 45 | 75 | 4 | 0 | 0 | 0 | 0 | 0 | 0 | 0 | 0 | 0 | 0 | 0 | 0 | 7 | 0 | 0 |
| 46 | 28 | 0 | 0 | 0 | 0 | 0 | 0 | 0 | 0 | 0 | 1 | 0 | 1 | 0 | 4 | 0 | 0 |
| 47 | 27 | 4 | 0 | 0 | 0 | 0 | 0 | 0 | 0 | 2 | 0 | 0 | 1 | 0 | 4 | 1 | 0 |
| 48 | 14 | 14 | 0 | 0 | 0 | 0 | 0 | 0 | 0 | 0 | 0 | 0 | 0 | 0 | 3 | 1 | 1 |
| 49 | 13 | 2 | 0 | 0 | 0 | 1 | 0 | 0 | 0 | 0 | 6 | 0 | 0 | 0 | 1 | 0 | 0 |
| 50 | 9 | 0 | 0 | 0 | 0 | 0 | 0 | 0 | 0 | 0 | 5 | 0 | 0 | 0 | 0 | 0 | 0 |
| 51 | 4 | 0 | 0 | 0 | 0 | 1 | 0 | 0 | 0 | 0 | 3 | 0 | 0 | 0 | 0 | 0 | 0 |
| 52 | 0 | 0 | 0 | 0 | 0 | 0 | 0 | 0 | 0 | 0 | 0 | 0 | 0 | 0 | 0 | 0 | 0 |
| 53 | 2 | 2 | 0 | 0 | 1 | 0 | 0 | 1 | 0 | 2 | 42 | 0 | 1 | 0 | 0 | 0 | 0 |
| 54 | 0 | 0 | 0 | 0 | 2 | 0 | 0 | 13 | 0 | 1 | 254 | 0 | 0 | 0 | 0 | 0 | 0 |
| 55 | 4 | 0 | 0 | 0 | 1 | 0 | 0 | 6 | 0 | 0 | 45 | 0 | 0 | 0 | 1 | 0 | 0 |
| 56 | 6 | 0 | 0 | 0 | 2 | 1 | 0 | 3 | 0 | 0 | 183 | 0 | 0 | 0 | 0 | 0 | 0 |
| 57 | 0 | 0 | 0 | 0 | 0 | 0 | 0 | 0 | 0 | 0 | 10 | 0 | 0 | 0 | 0 | 0 | 0 |
| 58 | 0 | 0 | 0 | 0 | 0 | 0 | 0 | 1 | 0 | 0 | 70 | 0 | 0 | 0 | 0 | 0 | 0 |
| 59 | 14 | 2 | 0 | 0 | 5 | 5 | 0 | 2 | 0 | 0 | 200 | 0 | 0 | 0 | 0 | 0 | 0 |
| 60 | 1 | 0 | 0 | 0 | 2 | 0 | 0 | 7 | 0 | 0 | 91 | 0 | 0 | 0 | 0 | 0 | 0 |
| 61 | 0 | 0 | 0 | 0 | 2 | 0 | 0 | 4 | 0 | 1 | 199 | 0 | 0 | 0 | 0 | 0 | 0 |
| 62 | 0 | 0 | 0 | 0 | 0 | 1 | 0 | 0 | 0 | 0 | 0 | 0 | 0 | 0 | 0 | 0 | 0 |
| 63 | 40 | 0 | 0 | 0 | 0 | 0 | 0 | 0 | 0 | 0 | 18 | 0 | 0 | 0 | 5 | 0 | 0 |
| 64 | 3 | 0 | 0 | 0 | 0 | 0 | 0 | 0 | 0 | 0 | 13 | 0 | 0 | 0 | 0 | 0 | 0 |
| 65 | 2 | 2 | 0 | 0 | 2 | 0 | 3 | 3 | 0 | 0 | 172 | 0 | 0 | 0 | 1 | 1 | 0 |
| 66 | 82 | 3 | 0 | 0 | 1 | 0 | 0 | 0 | 0 | 3 | 31 | 0 | 0 | 0 | 10 | 2 | 1 |
| 67 | 0 | 0 | 0 | 0 | 2 | 0 | 2 | 0 | 0 | 0 | 146 | 0 | 0 | 0 | 0 | 0 | 0 |
| 68 | 39 | 5 | 0 | 0 | 1 | 0 | 2 | 0 | 0 | 0 | 37 | 0 | 0 | 0 | 4 | 1 | 0 |
| 69 | 1 | 0 | 0 | 0 | 1 | 0 | 1 | 0 | 0 | 1 | 33 | 0 | 0 | 0 | 0 | 0 | 0 |
| 70 | 1 | 0 | 0 | 0 | 1 | 1 | 0 | 1 | 0 | 0 | 19 | 0 | 0 | 0 | 0 | 0 | 0 |
| 71 | 40 | 1 | 0 | 0 | 0 | 2 | 1 | 0 | 0 | 0 | 13 | 0 | 0 | 0 | 3 | 0 | 0 |
| 72 | 0 | 0 | 0 | 0 | 0 | 0 | 0 | 0 | 0 | 0 | 0 | 0 | 0 | 0 | 0 | 0 | 0 |
| 73 | 0 | 0 | 0 | 0 | 0 | 0 | 0 | 0 | 0 | 0 | 2 | 0 | 0 | 0 | 0 | 0 | 0 |
| 74 | 4 | 0 | 0 | 0 | 0 | 0 | 0 | 0 | 1 | 0 | 1 | 0 | 0 | 0 | 0 | 0 | 0 |
| 75 | 20 | 1 | 0 | 0 | 2 | 0 | 0 | 0 | 0 | 0 | 4 | 0 | 0 | 0 | 2 | 0 | 0 |
| 76 | 1 | 0 | 0 | 0 | 1 | 0 | 0 | 1 | 0 | 0 | 2 | 0 | 0 | 0 | 0 | 0 | 0 |
| 77 | 5 | 0 | 0 | 0 | 0 | 1 | 0 | 0 | 0 | 0 | 1 | 0 | 0 | 0 | 1 | 0 | 0 |
| 78 | 4 | 0 | 0 | 0 | 0 | 1 | 0 | 0 | 0 | 0 | 2 | 0 | 0 | 0 | 1 | 0 | 0 |
| 79 | 97 | 1 | 0 | 0 | 0 | 1 | 0 | 0 | 0 | 0 | 12 | 0 | 0 | 0 | 11 | 0 | 0 |
| 80 | 5 | 2 | 0 | 0 | 0 | 0 | 0 | 0 | 0 | 0 | 18 | 0 | 0 | 0 | 1 | 0 | 0 |
| 81 | 7 | 0 | 0 | 0 | 2 | 0 | 0 | 0 | 0 | 0 | 24 | 0 | 0 | 0 | 1 | 0 | 0 |
| 82 | 3 | 1 | 0 | 0 | 0 | 0 | 0 | 0 | 0 | 0 | 1 | 0 | 0 | 0 | 0 | 0 | 0 |
| 83 | 1 | 0 | 0 | 0 | 0 | 0 | 0 | 0 | 0 | 0 | 0 | 0 | 0 | 0 | 0 | 0 | 0 |
| 84 | 4 | 0 | 0 | 0 | 0 | 0 | 0 | 1 | 0 | 0 | 2 | 0 | 0 | 0 | 2 | 0 | 0 |
| 85 | 2 | 0 | 0 | 0 | 0 | 0 | 0 | 1 | 0 | 0 | 3 | 0 | 0 | 0 | 1 | 0 | 0 |
| 86 | 37 | 3 | 0 | 0 | 1 | 0 | 0 | 0 | 0 | 0 | 8 | 0 | 0 | 0 | 4 | 1 | 0 |
| 87 | 26 | 4 | 0 | 0 | 0 | 0 | 0 | 0 | 0 | 0 | 4 | 0 | 0 | 0 | 4 | 1 | 0 |
| 88 | 4 | 3 | 0 | 0 | 1 | 0 | 2 | 0 | 0 | 0 | 6 | 0 | 0 | 0 | 2 | 1 | 0 |
| 89 | 76 | 11 | 0 | 0 | 0 | 0 | 1 | 0 | 0 | 0 | 7 | 0 | 0 | 2 | 8 | 2 | 0 |
| 90 | 9 | 3 | 0 | 0 | 0 | 1 | 0 | 0 | 0 | 0 | 19 | 0 | 0 | 0 | 1 | 1 | 0 |
| 91 | 2 | 6 | 0 | 0 | 0 | 0 | 0 | 0 | 0 | 0 | 11 | 0 | 0 | 0 | 2 | 1 | 0 |
| 92 | 10 | 0 | 0 | 0 | 0 | 0 | 0 | 0 | 0 | 0 | 10 | 0 | 0 | 0 | 1 | 0 | 0 |
| 93 | 3 | 1 | 0 | 0 | 1 | 1 | 0 | 1 | 0 | 0 | 9 | 0 | 0 | 0 | 2 | 1 | 0 |
| 94 | 43 | 12 | 0 | 0 | 1 | 0 | 0 | 1 | 0 | 0 | 18 | 0 | 0 | 1 | 6 | 1 | 0 |
| 95 | 3 | 0 | 0 | 0 | 0 | 2 | 0 | 0 | 0 | 0 | 41 | 0 | 0 | 0 | 0 | 0 | 0 |
| 96 | 8 | 1 | 0 | 0 | 0 | 0 | 0 | 0 | 0 | 0 | 7 | 0 | 0 | 0 | 1 | 0 | 0 |
| 97 | 0 | 0 | 0 | 0 | 0 | 0 | 0 | 0 | 0 | 0 | 0 | 0 | 0 | 0 | 0 | 0 | 0 |
| 98 | 0 | 0 | 0 | 0 | 0 | 0 | 0 | 0 | 0 | 0 | 1 | 0 | 0 | 0 | 0 | 0 | 0 |
| 99 | 0 | 0 | 0 | 0 | 0 | 0 | 0 | 0 | 0 | 0 | 4 | 0 | 0 | 0 | 0 | 0 | 0 |
| 100 | 1 | 1 | 0 | 0 | 0 | 0 | 0 | 1 | 0 | 0 | 28 | 0 | 0 | 0 | 1 | 1 | 0 |
| 101 | 4 | 2 | 0 | 0 | 0 | 0 | 0 | 0 | 0 | 0 | 0 | 0 | 0 | 0 | 1 | 0 | 0 |
| 102 | 11 | 1 | 0 | 0 | 0 | 0 | 0 | 0 | 0 | 1 | 4 | 0 | 0 | 0 | 2 | 0 | 0 |
| 103 | 15 | 1 | 0 | 0 | 1 | 0 | 0 | 0 | 0 | 1 | 3 | 0 | 0 | 0 | 3 | 0 | 0 |
| 104 | 15 | 5 | 0 | 0 | 0 | 1 | 0 | 0 | 0 | 1 | 31 | 0 | 1 | 0 | 1 | 1 | 0 |
| 105 | 24 | 1 | 0 | 0 | 0 | 0 | 0 | 0 | 0 | 0 | 2 | 0 | 0 | 0 | 4 | 0 | 0 |
| 106 | 1 | 1 | 0 | 0 | 0 | 0 | 0 | 0 | 0 | 0 | 0 | 0 | 0 | 0 | 1 | 0 | 0 |
| 107 | 17 | 1 | 0 | 0 | 0 | 0 | 0 | 0 | 0 | 0 | 0 | 0 | 0 | 0 | 4 | 1 | 0 |
| 108 | 4 | 0 | 0 | 0 | 0 | 0 | 0 | 0 | 0 | 0 | 0 | 0 | 0 | 0 | 1 | 0 | 0 |
| 109 | 18 | 1 | 0 | 0 | 0 | 0 | 0 | 0 | 0 | 0 | 0 | 0 | 0 | 0 | 2 | 0 | 0 |
| 110 | 4 | 0 | 0 | 0 | 0 | 1 | 0 | 0 | 0 | 0 | 0 | 0 | 0 | 0 | 1 | 0 | 0 |
| 111 | 1 | 0 | 0 | 0 | 0 | 0 | 0 | 0 | 0 | 0 | 0 | 0 | 0 | 0 | 0 | 0 | 0 |
| 112 | 37 | 3 | 0 | 0 | 0 | 0 | 0 | 0 | 0 | 0 | 0 | 1 | 0 | 1 | 3 | 1 | 0 |
| 113 | 10 | 1 | 1 | 0 | 0 | 0 | 0 | 0 | 0 | 0 | 0 | 0 | 0 | 0 | 1 | 0 | 0 |
| 114 | 6 | 4 | 0 | 0 | 0 | 0 | 0 | 0 | 0 | 0 | 0 | 0 | 0 | 0 | 1 | 0 | 0 |
| 115 | 1 | 0 | 0 | 0 | 0 | 0 | 0 | 0 | 0 | 0 | 0 | 0 | 0 | 0 | 0 | 0 | 0 |
| 116 | 0 | 0 | 0 | 0 | 0 | 0 | 0 | 0 | 0 | 0 | 0 | 0 | 0 | 0 | 0 | 0 | 0 |
| 117 | 1 | 1 | 0 | 0 | 0 | 0 | 0 | 0 | 0 | 0 | 1 | 0 | 0 | 0 | 0 | 0 | 0 |
| 118 | 0 | 0 | 0 | 0 | 0 | 0 | 0 | 0 | 0 | 0 | 0 | 0 | 0 | 0 | 0 | 0 | 0 |
| 119 | 0 | 0 | 0 | 0 | 0 | 0 | 0 | 0 | 0 | 0 | 0 | 0 | 0 | 0 | 0 | 0 | 0 |
| 120 | 0 | 0 | 0 | 0 | 0 | 0 | 0 | 0 | 0 | 0 | 0 | 0 | 0 | 0 | 0 | 0 | 0 |
| 121 | 0 | 0 | 0 | 0 | 0 | 0 | 0 | 0 | 0 | 0 | 0 | 0 | 0 | 0 | 0 | 0 | 0 |
| 122 | 3 | 0 | 0 | 0 | 0 | 0 | 0 | 0 | 0 | 0 | 0 | 0 | 0 | 0 | 2 | 0 | 0 |
| 123 | 7 | 2 | 0 | 0 | 0 | 0 | 0 | 0 | 0 | 0 | 1 | 0 | 0 | 0 | 2 | 0 | 0 |
| 124 | 6 | 2 | 0 | 0 | 0 | 0 | 0 | 0 | 0 | 0 | 0 | 0 | 0 | 0 | 2 | 0 | 0 |
| 125 | 20 | 7 | 0 | 0 | 0 | 0 | 0 | 0 | 0 | 1 | 0 | 0 | 0 | 0 | 3 | 1 | 0 |
| 126 | 22 | 8 | 0 | 0 | 0 | 0 | 0 | 0 | 0 | 0 | 0 | 0 | 0 | 0 | 3 | 1 | 0 |
| 127 | 24 | 0 | 0 | 0 | 0 | 0 | 0 | 0 | 0 | 0 | 0 | 0 | 0 | 0 | 3 | 0 | 0 |
| 128 | 33 | 8 | 0 | 0 | 0 | 0 | 0 | 0 | 0 | 0 | 0 | 0 | 0 | 0 | 4 | 0 | 1 |
| 129 | 0 | 0 | 0 | 0 | 0 | 0 | 0 | 0 | 0 | 0 | 0 | 0 | 0 | 0 | 0 | 0 | 0 |
| 130 | 2 | 0 | 0 | 0 | 0 | 0 | 0 | 0 | 0 | 0 | 0 | 0 | 0 | 0 | 1 | 0 | 0 |
| 131 | 0 | 0 | 0 | 0 | 0 | 0 | 0 | 0 | 0 | 0 | 0 | 0 | 0 | 0 | 0 | 0 | 0 |
| 132 | 6 | 0 | 0 | 0 | 0 | 0 | 0 | 0 | 0 | 0 | 0 | 0 | 0 | 0 | 1 | 0 | 0 |
| 133 | 15 | 0 | 0 | 0 | 0 | 1 | 0 | 0 | 0 | 0 | 1 | 0 | 0 | 0 | 2 | 0 | 0 |
| 134 | 12 | 1 | 0 | 0 | 0 | 0 | 0 | 0 | 0 | 0 | 1 | 0 | 0 | 0 | 1 | 0 | 0 |
| 135 | 15 | 2 | 0 | 0 | 0 | 0 | 0 | 0 | 0 | 0 | 0 | 0 | 0 | 0 | 1 | 0 | 0 |
| 136 | 0 | 0 | 0 | 0 | 0 | 0 | 0 | 0 | 0 | 0 | 0 | 0 | 0 | 0 | 0 | 0 | 0 |
| 137 | 22 | 5 | 0 | 0 | 0 | 1 | 0 | 0 | 0 | 0 | 1 | 0 | 0 | 0 | 3 | 0 | 0 |
| 138 | 51 | 7 | 0 | 0 | 0 | 0 | 0 | 0 | 0 | 0 | 0 | 0 | 0 | 0 | 5 | 0 | 0 |
| 139 | 18 | 0 | 0 | 0 | 0 | 0 | 0 | 0 | 0 | 0 | 0 | 0 | 0 | 0 | 3 | 0 | 0 |
| 140 | 174 | 1 | 0 | 0 | 0 | 0 | 0 | 0 | 0 | 0 | 0 | 0 | 1 | 0 | 16 | 0 | 0 |
| 141 | 35 | 8 | 0 | 0 | 0 | 0 | 0 | 0 | 0 | 0 | 0 | 0 | 0 | 0 | 4 | 1 | 0 |
| 142 | 22 | 1 | 0 | 0 | 0 | 0 | 0 | 0 | 0 | 0 | 1 | 0 | 0 | 0 | 3 | 0 | 0 |
| 143 | 3 | 1 | 0 | 0 | 0 | 0 | 0 | 0 | 0 | 0 | 0 | 0 | 0 | 0 | 1 | 0 | 0 |
| 144 | 33 | 2 | 0 | 0 | 0 | 0 | 0 | 0 | 0 | 0 | 0 | 0 | 0 | 0 | 3 | 0 | 0 |
| 145 | 112 | 71 | 0 | 0 | 0 | 0 | 0 | 0 | 0 | 1 | 0 | 1 | 0 | 1 | 19 | 7 | 1 |
| 146 | 5 | 1 | 0 | 0 | 0 | 0 | 0 | 0 | 0 | 0 | 0 | 0 | 0 | 0 | 1 | 0 | 0 |
| 147 | 84 | 3 | 0 | 0 | 0 | 0 | 0 | 0 | 0 | 0 | 0 | 0 | 0 | 0 | 7 | 0 | 0 |
| 148 | 120 | 17 | 2 | 0 | 0 | 0 | 0 | 0 | 0 | 0 | 0 | 1 | 0 | 0 | 16 | 2 | 2 |
| 149 | 22 | 2 | 0 | 0 | 0 | 0 | 0 | 0 | 0 | 0 | 0 | 0 | 0 | 0 | 3 | 0 | 0 |
| 150 | 42 | 3 | 0 | 0 | 0 | 0 | 0 | 0 | 0 | 0 | 0 | 0 | 0 | 0 | 5 | 0 | 0 |
| 151 | 14 | 1 | 0 | 0 | 0 | 0 | 0 | 0 | 0 | 0 | 0 | 0 | 0 | 0 | 2 | 0 | 0 |
| 152 | 13 | 0 | 0 | 0 | 0 | 0 | 0 | 0 | 0 | 0 | 2 | 0 | 0 | 0 | 1 | 0 | 0 |
| 153 | 5 | 0 | 0 | 0 | 0 | 1 | 0 | 0 | 0 | 0 | 3 | 0 | 0 | 0 | 1 | 0 | 0 |
| 154 | 3 | 1 | 0 | 0 | 0 | 1 | 0 | 0 | 0 | 0 | 25 | 0 | 0 | 0 | 0 | 0 | 0 |
| 155 | 17 | 1 | 0 | 0 | 0 | 1 | 0 | 0 | 0 | 0 | 29 | 0 | 0 | 0 | 2 | 0 | 0 |
| 156 | 6 | 0 | 0 | 0 | 0 | 0 | 0 | 0 | 0 | 0 | 8 | 0 | 0 | 0 | 1 | 0 | 0 |
| 157 | 11 | 0 | 0 | 0 | 0 | 0 | 0 | 0 | 0 | 0 | 5 | 0 | 0 | 0 | 1 | 0 | 0 |
| 158 | 37 | 0 | 0 | 0 | 0 | 0 | 0 | 0 | 0 | 1 | 17 | 0 | 0 | 0 | 2 | 0 | 0 |
| 159 | 39 | 1 | 0 | 0 | 0 | 0 | 0 | 0 | 0 | 0 | 2 | 1 | 0 | 0 | 5 | 1 | 0 |
| 160 | 7 | 1 | 0 | 0 | 0 | 0 | 0 | 0 | 0 | 3 | 108 | 0 | 0 | 0 | 1 | 0 | 0 |
| 161 | 135 | 0 | 0 | 0 | 1 | 0 | 0 | 0 | 0 | 0 | 32 | 0 | 0 | 0 | 15 | 0 | 1 |
| 162 | 27 | 15 | 0 | 0 | 0 | 0 | 0 | 0 | 0 | 0 | 0 | 0 | 0 | 1 | 6 | 2 | 0 |
| 163 | 59 | 0 | 0 | 0 | 0 | 0 | 0 | 0 | 0 | 0 | 0 | 0 | 0 | 0 | 6 | 0 | 0 |
| 164 | 21 | 0 | 0 | 0 | 0 | 0 | 0 | 0 | 0 | 0 | 0 | 0 | 0 | 0 | 3 | 0 | 0 |
| 165 | 0 | 0 | 0 | 0 | 0 | 0 | 0 | 0 | 0 | 0 | 15 | 0 | 0 | 0 | 0 | 0 | 0 |
| 166 | 3 | 0 | 0 | 0 | 0 | 0 | 0 | 0 | 0 | 0 | 18 | 0 | 0 | 0 | 2 | 0 | 0 |
| 167 | 8 | 0 | 0 | 0 | 0 | 0 | 0 | 1 | 0 | 0 | 19 | 0 | 0 | 0 | 1 | 0 | 0 |
| 168 | 7 | 1 | 0 | 0 | 0 | 0 | 0 | 0 | 0 | 0 | 5 | 0 | 0 | 0 | 0 | 0 | 0 |
| 169 | 51 | 4 | 0 | 0 | 0 | 0 | 0 | 0 | 0 | 0 | 0 | 0 | 0 | 0 | 7 | 1 | 0 |
| 170 | 67 | 6 | 0 | 0 | 0 | 0 | 0 | 0 | 0 | 0 | 1 | 0 | 0 | 0 | 9 | 1 | 0 |
| 171 | 37 | 5 | 0 | 0 | 0 | 0 | 0 | 0 | 0 | 0 | 1 | 0 | 0 | 0 | 4 | 0 | 0 |
