## Supplemental Table S2 for "Spatial distribution of ticks and tick-borne pathogens in central Hokkaido, Japan and associated ecological factors revealed by intensive short-term survey in 2024"

|  | Species | Number of present sites |
| --- | --- | --- |
| Tick | *I. ovatus* | 139 |
|  | *I. persulcatus* | 97 |
|  | *I. pavlovskyi* | 10 |
|  | *H. megaspinosa* | 47 |
|  | *H. longicornis* | 22 |
|  | *H. flava* | 41 |
|  | *H. japonica* | 10 |
| Pathogen | TBEV | 7 |
|  | YEZV | 7 |
|  | BJNV | 9 |
|  | LDB | 117 |
|  | pLDB | 42 |
|  | RFB | 13 |
