## Supplemental Table S3 for "Spatial distribution of ticks and tick-borne pathogens in central Hokkaido, Japan and associated ecological factors revealed by intensive short-term survey in 2024"

| | | r = 0.1 | r = 0.2 | r = 0.3 | r = 0.4 | r = 0.5 | r = 0.6 | r = 0.7 | r = 0.8 | r = 0.9 | r = 1 | r = 2 | r = 3 | r = 4 | r = 5 | r = 6 | r = 7 | r = 8 | r = 9 | r = 10 | r = 11 | r = 12 | r = 13 | r = 14 | r = 15 | r = 16 | r = 17 | r = 18 | r = 19 | r = 20 |
|---|---|---|---|---|---|---|---|---|---|---|---|---|---|---|---|---|---|---|---|---|---|---|---|---|---|---|---|---|---|---|
| Iov | Moran's I | 0.38 | 0.35 | 0.34 | 0.32 | 0.31 | 0.30 | 0.30 | 0.29 | 0.29 | 0.28 | 0.26 | 0.24 | 0.23 | 0.21 | 0.19 | 0.18 | 0.16 | 0.15 | 0.14 | 0.12 | 0.12 | 0.11 | 0.10 | 0.09 | 0.09 | 0.08 | 0.08 | 0.07 | 0.07 |
| | p-value | <0.001 | <0.001 | <0.001 | <0.001 | <0.001 | <0.001 | <0.001 | <0.001 | <0.001 | <0.001 | <0.001 | <0.001 | <0.001 | <0.001 | <0.001 | <0.001 | <0.001 | <0.001 | <0.001 | <0.001 | <0.001 | <0.001 | <0.001 | <0.001 | <0.001 | <0.001 | <0.001 | <0.001 | <0.001 |
| Ipe | Moran's I | 0.15 | 0.15 | 0.15 | 0.15 | 0.16 | 0.16 | 0.16 | 0.16 | 0.16 | 0.17 | 0.17 | 0.16 | 0.15 | 0.14 | 0.13 | 0.12 | 0.11 | 0.10 | 0.09 | 0.08 | 0.08 | 0.07 | 0.07 | 0.06 | 0.06 | 0.05 | 0.05 | 0.05 | 0.04 |
| | p-value | 0.023 | 0.020 | 0.017 | 0.014 | 0.012 | 0.010 | 0.008 | 0.006 | 0.005 | 0.004 | <0.001 | <0.001 | <0.001 | <0.001 | <0.001 | <0.001 | <0.001 | <0.001 | <0.001 | <0.001 | <0.001 | <0.001 | <0.001 | <0.001 | <0.001 | <0.001 | <0.001 | <0.001 | <0.001 |
| Ipa | Moran's I | 0.23 | 0.22 | 0.22 | 0.21 | 0.21 | 0.21 | 0.21 | 0.21 | 0.21 | 0.21 | 0.20 | 0.18 | 0.15 | 0.13 | 0.12 | 0.10 | 0.09 | 0.08 | 0.07 | 0.06 | 0.06 | 0.05 | 0.05 | 0.04 | 0.04 | 0.04 | 0.03 | 0.03 | 0.03 |
| | p-value | 0.003 | 0.003 | 0.003 | 0.003 | 0.003 | 0.003 | 0.003 | 0.002 | 0.002 | 0.001 | <0.001 | <0.001 | <0.001 | <0.001 | <0.001 | <0.001 | <0.001 | <0.001 | <0.001 | <0.001 | <0.001 | <0.001 | <0.001 | <0.001 | <0.001 | <0.001 | <0.001 | <0.001 | <0.001 |
| Hme | Moran's I | 0.51 | 0.53 | 0.54 | 0.54 | 0.53 | 0.53 | 0.53 | 0.52 | 0.52 | 0.51 | 0.46 | 0.43 | 0.40 | 0.37 | 0.35 | 0.32 | 0.30 | 0.28 | 0.26 | 0.25 | 0.24 | 0.22 | 0.21 | 0.20 | 0.19 | 0.19 | 0.18 | 0.17 | 0.16 |
| | p-value | <0.001 | <0.001 | <0.001 | <0.001 | <0.001 | <0.001 | <0.001 | <0.001 | <0.001 | <0.001 | <0.001 | <0.001 | <0.001 | <0.001 | <0.001 | <0.001 | <0.001 | <0.001 | <0.001 | <0.001 | <0.001 | <0.001 | <0.001 | <0.001 | <0.001 | <0.001 | <0.001 | <0.001 | <0.001 |
| Hlo | Moran's I | 1.06 | 1.05 | 1.05 | 1.04 | 1.03 | 1.01 | 1.00 | 0.98 | 0.96 | 0.94 | 0.77 | 0.65 | 0.55 | 0.48 | 0.42 | 0.37 | 0.33 | 0.30 | 0.27 | 0.25 | 0.23 | 0.21 | 0.19 | 0.18 | 0.16 | 0.15 | 0.14 | 0.14 | 0.13 |
| | p-value | <0.001 | <0.001 | <0.001 | <0.001 | <0.001 | <0.001 | <0.001 | <0.001 | <0.001 | <0.001 | <0.001 | <0.001 | <0.001 | <0.001 | <0.001 | <0.001 | <0.001 | <0.001 | <0.001 | <0.001 | <0.001 | <0.001 | <0.001 | <0.001 | <0.001 | <0.001 | <0.001 | <0.001 | <0.001 |
| Hfl | Moran's I | 0.13 | 0.14 | 0.14 | 0.13 | 0.13 | 0.12 | 0.11 | 0.11 | 0.10 | 0.10 | 0.09 | 0.09 | 0.10 | 0.10 | 0.10 | 0.10 | 0.10 | 0.09 | 0.09 | 0.09 | 0.08 | 0.08 | 0.08 | 0.07 | 0.07 | 0.07 | 0.07 | 0.06 | 0.06 | 0.06 |
| | p-value | 0.062 | 0.057 | 0.056 | 0.059 | 0.063 | 0.069 | 0.074 | 0.078 | 0.081 | 0.083 | 0.056 | 0.019 | 0.005 | 0.001 | <0.001 | <0.001 | <0.001 | <0.001 | <0.001 | <0.001 | <0.001 | <0.001 | <0.001 | <0.001 | <0.001 | <0.001 | <0.001 | <0.001 | <0.001 |
| Hja | Moran's I | 0.31 | 0.34 | 0.36 | 0.37 | 0.38 | 0.38 | 0.39 | 0.39 | 0.39 | 0.38 | 0.31 | 0.23 | 0.18 | 0.14 | 0.12 | 0.10 | 0.08 | 0.07 | 0.06 | 0.06 | 0.05 | 0.04 | 0.04 | 0.04 | 0.03 | 0.03 | 0.03 | 0.03 | 0.02 |
| | p-value | <0.001 | <0.001 | <0.001 | <0.001 | <0.001 | <0.001 | <0.001 | <0.001 | <0.001 | <0.001 | <0.001 | <0.001 | <0.001 | <0.001 | <0.001 | <0.001 | <0.001 | <0.001 | <0.001 | <0.001 | <0.001 | <0.001 | <0.001 | <0.001 | <0.001 | <0.001 | <0.001 | <0.001 | <0.001 |
| TBEV | Moran's I | 0.14 | 0.16 | 0.17 | 0.17 | 0.17 | 0.17 | 0.17 | 0.17 | 0.16 | 0.16 | 0.10 | 0.07 | 0.05 | 0.04 | 0.03 | 0.02 | 0.02 | 0.01 | 0.01 | 0.01 | 0.00 | 0.00 | 0.00 | 0.00 | 0.00 | 0.00 | 0.00 | 0.00 | 0.00 |
| | p-value | 0.048 | 0.027 | 0.020 | 0.017 | 0.015 | 0.014 | 0.014 | 0.014 | 0.015 | 0.016 | 0.035 | 0.056 | 0.075 | 0.094 | 0.115 | 0.136 | 0.159 | 0.182 | 0.205 | 0.227 | 0.248 | 0.268 | 0.287 | 0.305 | 0.321 | 0.337 | 0.351 | 0.364 | 0.377 |
| YEZV | Moran's I | -0.06 | -0.06 | -0.06 | -0.06 | -0.06 | -0.06 | -0.05 | -0.05 | -0.05 | -0.05 | -0.04 | -0.03 | -0.02 | -0.02 | -0.01 | -0.01 | -0.01 | 0.00 | 0.00 | 0.00 | 0.00 | 0.00 | 0.00 | 0.00 | 0.00 | 0.00 | 0.00 | 0.00 | 0.00 |
| | p-value | 0.736 | 0.737 | 0.734 | 0.731 | 0.729 | 0.729 | 0.730 | 0.732 | 0.734 | 0.736 | 0.744 | 0.726 | 0.688 | 0.639 | 0.586 | 0.536 | 0.492 | 0.456 | 0.427 | 0.406 | 0.391 | 0.380 | 0.373 | 0.370 | 0.368 | 0.369 | 0.370 | 0.373 | 0.376 |
| BJNV | Moran's I | -0.05 | -0.03 | -0.02 | 0.00 | 0.01 | 0.02 | 0.03 | 0.03 | 0.04 | 0.04 | 0.05 | 0.05 | 0.05 | 0.05 | 0.05 | 0.05 | 0.05 | 0.04 | 0.04 | 0.04 | 0.03 | 0.03 | 0.03 | 0.03 | 0.02 | 0.02 | 0.02 | 0.02 | 0.02 |
| | p-value | 0.671 | 0.611 | 0.547 | 0.485 | 0.430 | 0.385 | 0.350 | 0.321 | 0.298 | 0.279 | 0.180 | 0.114 | 0.064 | 0.035 | 0.020 | 0.013 | 0.009 | 0.007 | 0.006 | 0.006 | 0.005 | 0.005 | 0.006 | 0.006 | 0.007 | 0.007 | 0.008 | 0.009 | 0.010 |
| LDB | Moran's I | 0.37 | 0.36 | 0.36 | 0.36 | 0.36 | 0.36 | 0.36 | 0.36 | 0.36 | 0.35 | 0.33 | 0.31 | 0.29 | 0.27 | 0.26 | 0.25 | 0.24 | 0.22 | 0.21 | 0.20 | 0.19 | 0.18 | 0.18 | 0.17 | 0.16 | 0.15 | 0.15 | 0.14 | 0.13 |
| | p-value | <0.001 | <0.001 | <0.001 | <0.001 | <0.001 | <0.001 | <0.001 | <0.001 | <0.001 | <0.001 | <0.001 | <0.001 | <0.001 | <0.001 | <0.001 | <0.001 | <0.001 | <0.001 | <0.001 | <0.001 | <0.001 | <0.001 | <0.001 | <0.001 | <0.001 | <0.001 | <0.001 | <0.001 | <0.001 |
| pLDB | Moran's I | 0.42 | 0.43 | 0.43 | 0.43 | 0.43 | 0.42 | 0.42 | 0.41 | 0.41 | 0.40 | 0.35 | 0.30 | 0.26 | 0.22 | 0.19 | 0.17 | 0.15 | 0.14 | 0.12 | 0.11 | 0.10 | 0.09 | 0.08 | 0.08 | 0.07 | 0.06 | 0.06 | 0.06 | 0.05 |
| | p-value | <0.001 | <0.001 | <0.001 | <0.001 | <0.001 | <0.001 | <0.001 | <0.001 | <0.001 | <0.001 | <0.001 | <0.001 | <0.001 | <0.001 | <0.001 | <0.001 | <0.001 | <0.001 | <0.001 | <0.001 | <0.001 | <0.001 | <0.001 | <0.001 | <0.001 | <0.001 | <0.001 | <0.001 | <0.001 |
| RFB | Moran's I | 0.25 | 0.24 | 0.23 | 0.22 | 0.22 | 0.22 | 0.21 | 0.21 | 0.21 | 0.20 | 0.17 | 0.12 | 0.09 | 0.06 | 0.05 | 0.04 | 0.03 | 0.03 | 0.03 | 0.02 | 0.02 | 0.02 | 0.02 | 0.02 | 0.02 | 0.02 | 0.02 | 0.01 | 0.01 |
| | p-value | 0.003 | 0.003 | 0.004 | 0.004 | 0.005 | 0.005 | 0.004 | 0.004 | 0.004 | 0.003 | 0.002 | 0.005 | 0.010 | 0.018 | 0.027 | 0.035 | 0.039 | 0.041 | 0.041 | 0.040 | 0.038 | 0.035 | 0.033 | 0.031 | 0.029 | 0.027 | 0.025 | 0.024 | 0.023 |
