## Supplemental Table S4 for "Spatial distribution of ticks and tick-borne pathogens in central Hokkaido, Japan and associated ecological factors revealed by intensive short-term survey in 2024"

|  |  |  | LOOCV | |  | Moran’s *I* test for residuals | |
| --- | --- | --- | --- | --- | --- | --- | --- |
|  | Species |  | AUC | TSS |  | Moran’s *I* | p-value |
| Tick | *I. ovatus* |  | 0.88 | 0.64 |  | -0.02 | 0.585 |
|  | *I. persulcatus* |  | 0.84 | 0.53 |  | 0.07 | 0.187 |
|  | *I. pavlovskyi* |  | 0.82 | 0.66 |  | -0.18 | 0.988 |
|  | *H. megaspinosa* |  | 0.90 | 0.71 |  | -0.01 | 0.496 |
|  | *H. longicornis* |  | 0.86 | 0.63 |  | -0.05 | 0.700 |
|  | *H. flava* |  | 0.71 | 0.41 |  | -0.02 | 0.586 |
|  | *H. japonica* |  | 0.80 | 0.58 |  | -0.02 | 0.558 |
| Pathogen | TBEV |  | 0.75 | 0.53 |  | 0.10 | 0.133 |
|  | YEZV |  | 0.72 | 0.52 |  | -0.02 | 0.578 |
|  | BJNV |  | 0.87 | 0.69 |  | -0.12 | 0.934 |
|  | LDB |  | 0.83 | 0.57 |  | 0.07 | 0.182 |
|  | pLDB |  | 0.86 | 0.59 |  | -0.02 | 0.548 |
|  | RFB |  | 0.76 | 0.61 |  | 0.11 | 0.068 |
