## Supplemental Table S5 for "Spatial distribution of ticks and tick-borne pathogens in central Hokkaido, Japan and associated ecological factors revealed by intensive short-term survey in 2024"

|  | Response variable | Explanatory variable | Coefficient | SE | z-value | p-value |
| --- | --- | --- | --- | --- | --- | --- |
| Tick | *I. ovatus* | (Intercept) | 3.072 | 0.563 | 5.458 | <0.001 |
|  |  | *forest* | 0.448 | 0.389 | 1.151 | 0.250 |
|  |  | *snow* | 1.761 | 0.605 | 2.912 | 0.004 |
|  |  | *cold* | -0.713 | 0.344 | -2.071 | 0.038 |
|  |  | *angl* | 1.586 | 0.724 | 2.191 | 0.028 |
|  | *I. persulcatus* | (Intercept) | 0.885 | 0.449 | 1.970 | 0.049 |
|  |  | *forest* | 1.174 | 0.410 | 2.863 | 0.004 |
|  |  | *snow* | 0.317 | 0.376 | 0.844 | 0.398 |
|  |  | *prec6* | -0.959 | 0.287 | -3.348 | 0.001 |
|  |  | *cold* | -0.455 | 0.252 | -1.803 | 0.071 |
|  |  | *elev* | 2.425 | 0.845 | 2.871 | 0.004 |
|  | *I. pavlovskyi* | (Intercept) | -5.959 | 1.720 | -3.464 | <0.001 |
|  |  | *grass* | -3.697 | 1.914 | -1.931 | 0.053 |
|  |  | *elev* | 1.876 | 1.377 | 1.362 | 0.173 |
|  |  | ti (*lon, lat*) |  |  | 13.94 | 0.259 |
|  | *H. megaspinosa* | (Intercept) | -2.306 | 0.536 | -4.303 | <0.001 |
|  |  | *grass* | -1.016 | 0.450 | -2.255 | 0.024 |
|  |  | *snow* | -0.451 | 0.774 | -0.582 | 0.561 |
|  |  | s (*lon*) |  |  | 16.18 | 0.006 |
|  |  | s (*lat*) |  |  | 11.36 | <0.001 |
|  |  | ti (*lon, lat*) |  |  | 12.78 | 0.059 |
|  | *H. longicornis* | (Intercept) | -6.144 | 2.289 | -2.684 | 0.007 |
|  |  | *snow* | -5.880 | 2.636 | -2.231 | 0.026 |
|  |  | *temp6* | -3.040 | 2.036 | -1.493 | 0.135 |
|  |  | s (*lon*) |  |  | 2.246 | 0.369 |
|  |  | s (*lat*) |  |  | 0.538 | 0.772 |
|  |  | ti (*lon, lat*) |  |  | 12.956 | 0.133 |
|  | *H. flava* | (Intercept) | -1.768 | 0.324 | -5.457 | <0.001 |
|  |  | *forest* | 0.814 | 0.408 | 1.995 | 0.046 |
|  |  | *snow* | -1.158 | 0.526 | -2.203 | 0.028 |
|  |  | *angl* | 0.432 | 0.397 | 1.087 | 0.277 |
|  |  | s (*lon*) |  |  | 9.427 | 0.044 |
|  |  | s (*lat*) |  |  | 3.183 | 0.361 |
|  |  | ti (*lon, lat*) |  |  | 9.151 | 0.117 |
|  | *H. japonica* | (Intercept) | -17.590 | 19.929 | -0.883 | 0.377 |
|  |  | *snow* | -2.250 | 1.132 | -1.987 | 0.047 |
|  |  | *elev* | 6.667 | 2.286 | 2.916 | 0.004 |
|  |  | s (*lon*) |  |  | 5.252 | 0.688 |
| Pathogen | TBEV | (Intercept) | -1.622 | 0.818 | -1.983 | 0.047 |
|  |  | *forest* | -0.872 | 0.945 | -0.922 | 0.356 |
|  |  | *prec6* ^2 | -0.563 | 0.713 | -0.789 | 0.430 |
|  |  | *temp6* | -3.541 | 1.214 | -2.917 | 0.004 |
|  |  | *cold* | -0.585 | 0.604 | -0.969 | 0.333 |
|  |  | *elev* ^2 | -1.786 | 1.001 | -1.784 | 0.074 |
|  | YEZV | (Intercept) | -3.912 | 1.711 | -2.286 | 0.022 |
|  |  | *forest* | 3.045 | 2.117 | 1.439 | 0.150 |
|  |  | *prec6* | -0.445 | 0.515 | -0.863 | 0.388 |
|  |  | *angl* ^2 | -0.791 | 0.646 | -1.225 | 0.220 |
|  |  | *raccoon* | -1.907 | 0.889 | -2.144 | 0.032 |
|  | BJNV | (Intercept) | -2.458 | 0.640 | -3.839 | <0.001 |
|  |  | *cold* | -0.747 | 0.622 | -1.201 | 0.230 |
|  |  | *elev* | 1.473 | 0.505 | 2.916 | 0.004 |
|  |  | *tanuki* | -0.901 | 0.776 | -1.162 | 0.245 |
|  | LDB | (Intercept) | 1.839 | 0.475 | 3.874 | <0.001 |
|  |  | *forest* | 0.535 | 0.322 | 1.663 | 0.096 |
|  |  | *snow* | 1.492 | 0.409 | 3.646 | <0.001 |
|  |  | *cold* | -0.531 | 0.265 | -2.003 | 0.045 |
|  |  | *elev* | 1.416 | 0.798 | 1.775 | 0.076 |
|  | pLDB | (Intercept) | -1.836 | 0.704 | -2.610 | 0.009 |
|  |  | *grass* | -0.854 | 0.653 | -1.309 | 0.191 |
|  |  | *temp6* | -3.126 | 0.835 | -3.742 | <0.001 |
|  |  | s (*lon*) | 2.003 | 2.474 | 8.988 | 0.013 |
|  |  | s (*lat*) |  |  | 12.685 | 0.053 |
|  |  | ti (*lon, lat*) |  |  | 18.412 | 0.020 |
|  | RFB | (Intercept) | -3.598 | 0.809 | -4.446 | <0.001 |
|  |  | *prec6* | -0.030 | 0.552 | -0.054 | 0.957 |
|  |  | *cold* | -1.825 | 0.690 | -2.645 | 0.008 |
|  |  | s (*lon*) |  |  | 6.928 | 0.008 |
|  |  | ti (*lon, lat*) |  |  | 11.942 | 0.252 |
